# Identification of a bile acid-binding transcription factor in *Clostridioides difficile* using chemical proteomics

**DOI:** 10.1101/2022.05.26.493666

**Authors:** ER Forster, X Yang, HC Hang, A Shen

## Abstract

*Clostridioides difficile* is a Gram-positive anaerobic bacterium that is the leading cause of hospital-acquired gastroenteritis in the US. In the gut milieu, *C. difficile* encounters microbiota-derived bile acids capable of inhibiting its growth, which are thought to be a mechanism of colonization resistance. While the levels of certain bile acids in the gut correlate with susceptibility to *C. difficile* infection, their molecular targets in *C. difficile* remain unknown. In this study, we sought to use chemical proteomics to identify bile acid-interacting proteins in *C. difficile*. Using photoaffinity bile acid probes and chemical proteomics, we identified a previously uncharacterized MerR family protein, CD3583 (now BapR), as a putative bile acid-sensing transcription regulator. Our data indicate that BapR binds and is stabilized by lithocholic acid (LCA) in *C. difficile*. Although loss of BapR did not affect *C. difficile*’s sensitivity to LCA, Δ*bapR* cells elongated more in the presence of LCA compared to wild-type cells. Transcriptomics revealed that BapR regulates the expression of the gene clusters *mdeA-cd3573* and *cd0618-cd0616*, and *cwpV*, with the expression of the *mdeA-cd3573* locus being specifically de-repressed in the presence of LCA in a BapR-dependent manner. Electrophoretic mobility shift assays revealed that BapR directly binds to the *mdeA* promoter region. Since *mdeA* is involved in amino acid-related sulfur metabolism and the *mdeA-cd3573* locus encodes putative transporters, we propose that BapR senses a gastrointestinal tract-specific small molecule, LCA, as an environmental cue for metabolic adaptation.

## Introduction

*Clostridioides difficile* is a Gram-positive, anerobic, spore-forming bacterium that is a leading cause of nosocomial gastroenteritis worldwide (1, 2). In 2017, *C. difficile* was responsible for ∼460,000 infections and ∼13,000 deaths in the United States alone (3). *C. difficile* infections (CDIs) are transmitted by its aerotolerant spores, which germinate in the distal small intestine in response to cholate-derived bile acids, giving rise to toxin-producing vegetative cells that colonize the colon (4). While prior antibiotic use and the consequent disruption of the gut microbiota is a primary risk factor for CDI, the specific mechanisms by which the microbiota protect against CDI remain largely unknown. However, production of secondary bile acids by the gut microbiota has been proposed to be a major contributor to colonization resistance. Bacteria that produce secondary bile acids are sufficient to confer colonization resistance against CDI, and the levels of these secondary bile acids in the gut correlate with resistance to CDI (4–8). Additionally, *C. difficile* is highly sensitive to growth inhibition by secondary bile acids, with lithocholic acid (LCA) and its derivatives being the most potent inhibitors (6, 7, 9).

Bile acids are amphipathic, detergent-like molecules produced by the liver and secreted into the small intestine to aid in fat digestion. Host-derived primary bile acids are largely reabsorbed via enterohepatic recirculation in the distal small intestine, but approximately 5% reach the large intestine where they are transformed into secondary bile acids by the microbiota (10). In cecal samples obtained from individuals who died an unnatural death, total bile acids were measured at ∼200 - 1,000 μM (11). This pool consisted mostly of the secondary bile acids LCA and deoxycholic acid (DCA): on average, each represented ∼25-33% of the bile acids in the non-CDI cecum (11) and feces (10). Growing evidence suggests that microbiota-derived secondary bile acids have broad pathophysiological implications in the host from carcinogenesis (12) to immune modulation and inflammation (13–20) to pathogen resistance (9, 21, 22), including *C. difficile*. Indeed, bile acids cause a broad range of stresses in bacteria including membrane disruption, protein denaturation, iron and calcium chelation, and DNA damage (23, 24).

DCA and LCA are made from the primary bile acids cholic acid (CA) and chenodeoxycholic acid (CDCA), respectively, via 7α-dehydroxylation by a select few members of the gut microbiota (25–31). This activity is encoded by the bile acid-inducible (*bai*) operon (25–31), which is highly correlated with resistance to CDI in animal models (7, 30, 31). A few *bai* operon-positive *Clostridium* species, namely *C. scindens*, are each sufficient to protect mice against CDI (7, 32, 33), although *bai* operon mutant strains have not yet been tested presumably due to genetic limitations. Notably, sequestration of bile from the cecal contents of *C. scindens*-colonized mice with cholestyramine is sufficient to rescue *C. difficile* growth *ex vivo*, suggesting that the bile acids produced by *C. scindens* inhibit *C. difficile* infection (7). Further, DCA and LCA are undetectable in the cecal contents of antibiotic-treated mice and recurrent CDI patient feces, but they are present in gut environments that are inhibitory to *C. difficile* (4–6, 8, 34). Underscoring these correlations, concentrations of DCA and LCA present in CDI-resistant mice are sufficient to inhibit *C. difficile* growth *in vitro* (6), and LCA tolerance correlates with virulence across *C. difficile* strains in a murine model of infection (35). While subsequent studies suggest that 7α-dehydroxylating bacteria also inhibit *C. difficile* through bile acid-independent mechanisms such as antibiotic production and competition for Stickland metabolism substrates (32, 36), secondary bile acids nonetheless appear to be a major contributor.

Gut commensals and several gut pathogens have evolved mechanisms to resist or tolerate the toxicity of bile acids (23, 24, 37), yet resistance mechanisms remain undefined in *C. difficile*. Currently, little is known about *C. difficile*’s interactions with bile acids at the molecular level, although bile acids have been shown to modulate several aspects of *C. difficile* physiology. LCA causes loss of flagella and cell elongation (38) whereas DCA induces biofilm formation and represses sporulation (39). Toxin expression and/or activity is inhibited by LCA, DCA, isolithocholic acid (isoLCA), isodeoxycholic acid, and ursodeoxycholic acid, and the TcdB toxin was recently shown to bind several bile acids (39–41). This latter observation represents the sole biochemical evidence of a *C. difficile* protein specifically binding to a bile acid to date. Indeed, while genetic studies suggest that the CspC pseudoprotease is a receptor for the primary bile acid germinant, taurocholic acid (TCA) (42), direct binding of TCA or other cholate-derived bile acid germinants (43) to *C. difficile* spore proteins has not been demonstrated. Furthermore, no targets of bile acids in *C. difficile* vegetative cells have been identified.

In this study, we employed recently developed photoaffinity bile acid probes in a chemical proteomics screen to identify the targets of toxic bile acids in vegetative *C. difficile*. While our screen identified several essential proteins as targets of LCA-derived probes, a putative MerR family transcription factor, CD3583 (now BapR), was highly enriched. Here, we demonstrate that BapR binds to LCA in *C. difficile*. BapR appears to modulate cell length in response to LCA and directly represses the expression of a locus encoding genes involved in sulfur metabolism. Our data also indicate that BapR indirectly represses the expression of two additional loci encoding two other putative transcription factors and the cell wall protein CwpV.

## Results

### Bile acid probes label distinct *C. difficile* proteins in a dose-dependent manner

Coupling photoaffinity probes of microbiota metabolites with chemical proteomics affords new opportunities to characterize specific protein targets and elucidate small molecule mechanisms of action (44). For example, bile acid photoaffinity probes recently identified the HilD regulator of SPI-1 virulence gene expression as a target of CDCA in *Salmonella* Typhimurium (Yang X, Stein K, Hang HC in revision). Using a similar approach, we sought to uncover the targets of bile acids in *C. difficile* to gain insight into the mechanism of inhibition by DCA or LCA and potential adaptation of *C. difficile* to these toxic, microbiota-derived molecules. These probes are generated from naturally occurring bile acids with the addition of two functional groups for photocrosslinking and bioorthogonal detection (**Figure 1A**). The diazirine functional group allows for covalent crosslinking to interacting proteins using UV light, while the terminal alkyne tag allows for detection or isolation of interacting proteins after bioorthogonal labeling with fluorophore-azide or biotin-azide reagents, respectively (**Figure 1B**).

**Figure 1.**
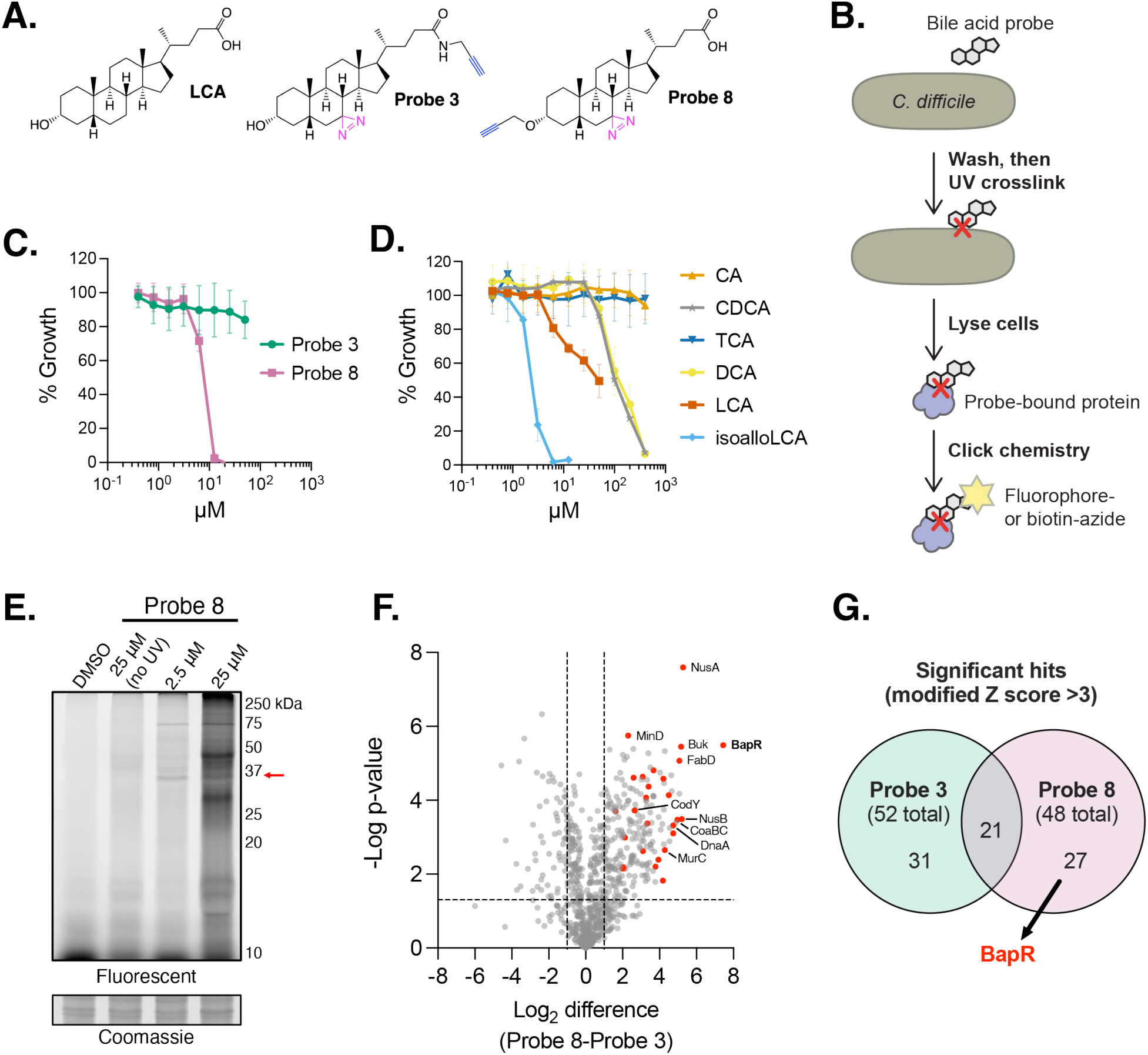
Identification of bile acid binding proteins by chemical proteomics. (**A**) Structures of bile acid probes; diazirine ring for UV crosslinking shown in magenta and terminal alkyne for click chemistry shown in blue. (**B**) Schematic of bile acid probe labeling and detection. (**C**) Inhibition of *C. difficile* growth in broth culture across various concentrations of bile acid probes calculated from OD_600_ after 5 hours of growth; error bars represent SD, n = 4 in two experiments. (**D**) Growth inhibition as in (C) with bile acids; error bars represent SD, n = 4 in two experiments. (**E**) In-gel fluorescent detection of probe labeling as illustrated in (C); cells were treated with probe for 30 minutes during log phase and Coomassie stain serves as a loading control, the gel is representative of three experiments. The red arrow indicates a band approximately the size of BapR. (**F**) Comparison of proteins isolated from *C. difficile* using Probe 8 vs Probe 3 and identified by LC-MS/MS; cells were grown with 10 μM probes for 1 hour during log-phase, dashed lines indicate p < 0.05 or 2-fold LFQ intensity difference and significant hits for Probe 8 (modified Z score > 3) are shown in red, n = 3. (**G**) Comparison of hits in each dataset, BapR is a significant hit for Probe 8 but not Probe 3.

To gain insight into how toxic bile acids affect *C. difficile*’s physiology, we sought to identify targets of these microbiota-derived molecules. Since LCA potently inhibits growth of *C. difficile*, we employed LCA-derived probes to identify LCA’s cellular targets. Using growth inhibition in broth culture as a readout of probe activity, we found that Probe 8 potently inhibited growth with an IC_50_ of ∼10 μM (**Figure 1C**). This inhibition was comparable to the IC_50_ of isoallolithocholic acid (isoalloLCA, ∼2 μM, **Figure 1D**), which was recently identified as an especially toxic bile acid to *C. difficile* in the gut of centenarians (9). Although solubility limited the range of concentrations we could test, Probe 8 was more potent than LCA (IC_50_ ∼50 μM) and DCA and CDCA (IC_50_s of ∼100 μM, **Figure 1D** **and Figure S2**). The primary bile acids CA and TCA did not inhibit growth up to 400 μM (**Figure 1D** **and Figure S2**). Importantly, we observed these effects at physiologically relevant concentrations since LCA and DCA have been measured at 1 - 450 μM and 30 - 700 μM, respectively, in non-CDI human cecal contents (11) and ∼1 mM and ∼1.2 mM, respectively, in human feces post-FMT (34). Interestingly, Probe 3 did not appreciably inhibit the growth of *C. difficile* (**Figure 1C**) despite sharing LCA as a parent structure with Probe 8. Due to its structural similarity to Probe 8 yet substantially reduced growth inhibitory activity, we selected Probe 3 as a control probe.

To qualitatively assess whether the probes specifically interact with proteins in *C. difficile*, we used fluorescent SDS-PAGE to visualize labeling. To this end, we treated log-phase *C. difficile* cultures with 2.5 or 25 μM Probe 8 or DMSO vehicle for 30 minutes, washed cells to remove unbound probe, and UV irradiated cells to covalently link the probe to interacting proteins. After generating total cell lysate and performing a click reaction in the total lysate to conjugate AlexaFluor-488-azide to the probe, these probe-bound proteins were visualized by SDS-PAGE using fluorescent imaging (**Figure 1B**). We observed distinct bands in a dose- and UV-dependent manner indicative of Probe 8 interacting with specific *C. difficile* proteins (**Figure 1E**). Possibly in line with its reduced growth inhibitory activity, Probe 3 exhibited weaker labeling (**Figure S1**).

### Chemical proteomics identifies bile acid-binding proteins in *C. difficile*

To identify the direct targets of Probe 8 visualized in the in-gel fluorescence assay, we used a chemical proteomics approach to enrich for probe-bound proteins. After conjugating biotin-azide onto probe-bound proteins (**Figure 1B**), we used streptavidin beads to isolate the proteins that interacted with Probe 8 and/or Probe 3 after 1 hour of treatment during log-phase (**Figure S1**). Streptavidin-enriched proteins were then identified using label-free quantitative liquid chromatography-tandem mass spectrometry (**Figure 1F** **and Table S1**). These proteomic analyses revealed 48 hits for Probe 8 and 52 hits for Probe 3 relative to DMSO (modified Z score > 3). Of the Probe 8 hits, 27 were specific to Probe 8 over Probe 3 (**Figure 1G**). We were particularly interested in Probe 8-specific proteins because Probe 8 is associated with growth inhibitory activity. Consistent with the toxicity of Probe 8 and its parent LCA molecule, 9 of the 27 hits specific to Probe 8 were essential (45). These included proteins involved in DNA replication, transcription, or cell division, namely DnaA, NusA, and MinD. The coenzyme A synthesis protein CoaBC and fatty acid synthesis protein FabD were additional essential hits. The essential cell wall synthesis protein MurC was of particular interest considering that *C. difficile* cells elongate with LCA treatment (38). However, the most enriched protein from our screen was a non-essential MerR family transcription factor, CD3583 (herein <u>b</u>ile <u>a</u>cid-binding <u>p</u>rotein regulator BapR; locus tag CD630_35830) (**Figure 1F**). MerR family transcription factors typically regulate efflux pumps that export the toxic molecules they sense (46) and can function as repressors in the absence of ligand as well as activators upon ligand binding. Given the role of MerR family transcription factors in regulating resistance or adaptative responses, we sought to explore the function of BapR in *C. difficile*.

### BapR specifically binds LCA-derived bile acids

Structural modeling using iTASSER (47) predicted that BapR is highly similar to a MerR family transcription factor in *Bacillus subtilis*, BmrR. BapR shares 18% identity and 28% similarity with BmrR, most of which is in the N-terminal region. MerR family transcription factors have conserved N-terminal DNA binding domains and variable C-terminal ligand-binding domains. These ligand binding domains have been observed to sense and respond to a wide diversity of molecules, with ligands ranging from metal ions to large amphipathic drugs (48). Although BmrR does not bind bile acids, its ligands are large hydrophobic molecules with some similarity to the sterol center of bile acids (49–51).

To validate BapR as a target of Probe 8, we used an independent method for assessing whether the bile acid probes could pull down BapR from *C. difficile*. Specifically, we treated *C. difficile* cultures expressing a FLAG-tagged allele of *bapR* from its native locus with either DMSO, Probe 8, or Probe 3 for 1 hour during log-phase. The probe was UV crosslinked to its protein targets, and then the cells were lysed. After conjugating biotin-azide to probe-bound proteins using click chemistry, we isolated these proteins using streptavidin beads (**Figure 1B**). FLAG-tagged BapR was enriched in the pull-downs using Probe 8, and to a lesser extent using Probe 3, relative to DMSO-treated cultures (**Figure 2A**, representative of three biological replicates). More biotinylated probe-bound protein was present in the Probe 8 pull-downs relative to Probe 3 as detected with fluorescent streptavidin, consistent with Probe 8 interacting with more proteins in our proteomics screen (**Figure 1F** **and Table S1**).

**Figure 2.**
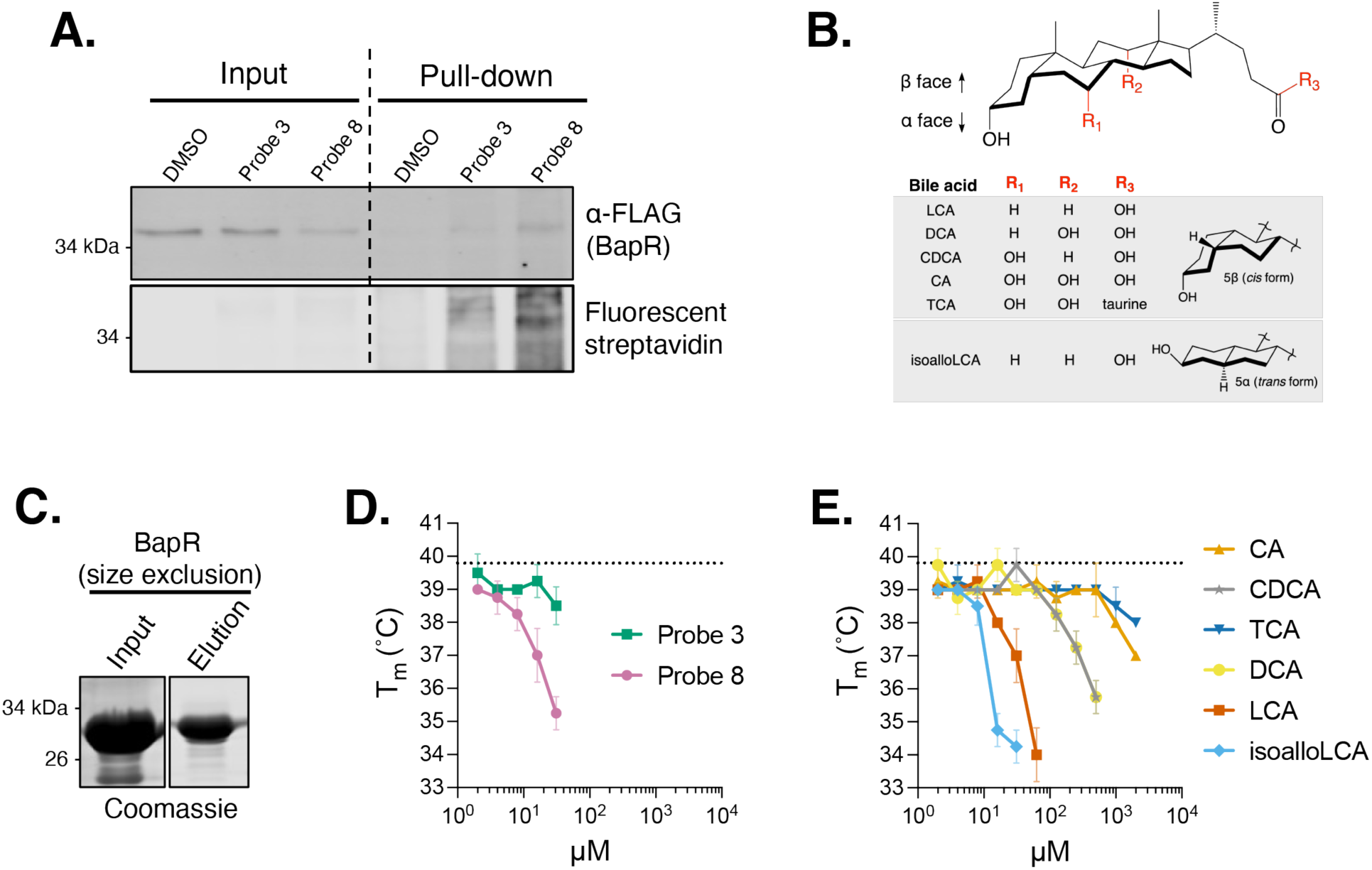
Validation of BapR as a bile acid-binding protein. (**A**) Pull-down of BapR from *C. difficile* using bile acid probes; cells were treated with 10 μM probes for 1 hour during log-phase before following the workflow in Figure 1C with biotin, then streptavidin beads pulled down the probe and BapR if bound. BapR was FLAG-tagged to facilitate its detection. Input samples were taken after the click reaction conjugating biotin to the probe, and fluorescent streptavidin detects presence of the probe. Blot is representative of 3 independent experiments. (**B**) Summarized structures of the bile acids used in this study. (**C**) Size exclusion chromatography of BapR after affinity purification. The gel is representative of 2 independent purifications. (**D**) Thermal shift assay with purified BapR; the melting temperature (T_m_) of BapR was assessed using SYPRO Orange dye across a range of bile acid probe concentrations; a change in melting temperature is indicative of binding. Dashed line indicates T_m_ of BapR in the presence of DMSO vehicle, error bars represent SD, n = 4 with protein from two independent protein purifications. (**E**) Thermal shift assay as in (D) with bile acids.

To determine whether we could directly detect binding between BapR and LCA, we used thermal shift assays to compare the relative affinity of BapR for different bile acids. Thermal shift assays (also known as differential scanning fluorimetry, DSF) measure the change in melting temperature (T_m_) of a protein in the presence of a ligand using a fluorescent dye that binds to hydrophobic regions of a protein as it unfolds. While ligand binding typically increases the T_m_ of a protein because ligand binding stabilizes proteins, it can in some instances decrease the T_m_ if the conformation stabilized by the ligand melts more readily (52). An example of this phenomenon can be found in isoLCA binding to the eukaryotic AKR1A1 aldo-keto reductase, for which isoLCA is a known ligand (53). Like AKR1A1, we observed a dose-dependent decrease in T_m_ when purified BapR (**Figure 2C**) was incubated with increasing concentrations of Probe 8 and certain bile acids in the presence of SYPRO Orange dye (**Figure 2D and E**). Specifically, we detected a ∼4°C T_m_ shift with 32 μM Probe 8 compared to a ∼1°C T_m_ shift with Probe 3 at the same concentration. We also observed up to a ∼5°C T_m_ shift at 64 μM LCA, whereas DCA and CDCA required ∼10-fold higher concentrations (500 µM) to shift the T_m_ of BapR by ∼3°C. Furthermore, CA and TCA were only able to shift the T_m_ by ∼1-2°C at 2 mM. It should be noted that the maximum concentration shown for each ligand tested was limited by measuring the non-specific interactions of a given bile acid with the SYPRO Orange dye (i.e. in the absence of BapR).

The apparent affinity of BapR for specific bile acids (LCA > DCA/CDCA > CA/TCA) correlates with fewer hydroxyl groups on the α face of the steroid center (**Figure 2B**), suggesting that hydrophobicity of the bile acid influences it interaction with BapR. IsoalloLCA differs from LCA in the orientation of the 3-OH and most notably by the stereochemistry at carbon 5: the 5β hydrogen in LCA results in a “bent” steroid ring, whereas the 5α hydrogen in isoalloLCA results in a more planar conformation. Interestingly, BapR appeared to have higher affinity for isoalloLCA given that it induced a ∼5°C T_m_ shift at a 2-fold lower concentration than LCA (32 µM vs. 64 μM, **Figure 2E**). The hydroxyl at carbon 3 is on the β face of the steroid center of isoalloLCA opposed to the α face for LCA, further implying that hydrophobicity of the α face is important for interaction with BapR.

### BapR influences cell length and interacts with LCA in *C. difficile*

After confirming that BapR binds bile acids, we sought to determine the physiological function of BapR in *C. difficile*. We initially hypothesized that BapR regulates resistance to bile acids like other MerR family transcription factors that regulate resistance to their ligands (48). However, we were unable to detect a growth defect for *C. difficile* lacking *bapR* in the presence of LCA (**Figure 3A**) or isoalloLCA (**Figure S3**). Besides growth inhibition, cell elongation and loss of flagella are the only known effects of LCA on *C. difficile* (38), so we instead asked whether BapR influences these phenotypes. While BapR did not influence motility on soft agar (data not shown), we observed that loss of BapR resulted in cells elongating ∼25% more than wild-type (WT) cells after a 3-hour exposure to 20 μM LCA (**Figure 3B**). Specifically, although LCA treatment increased the length of WT *C. difficile* cells by ∼2 μm, Δ*bapR* cells elongated an additional ∼0.5 μm over the WT strain under these conditions (**Figure 3C**). This phenotype was complemented by expressing *bapR* from its native promoter at the ectopic *pyrE* locus on the chromosome (Δ*bapR*/*bapR*; **Figure 3C**) (54). We did not observe gross morphological differences aside from cell length between the strains upon LCA exposure.

**Figure 3.**
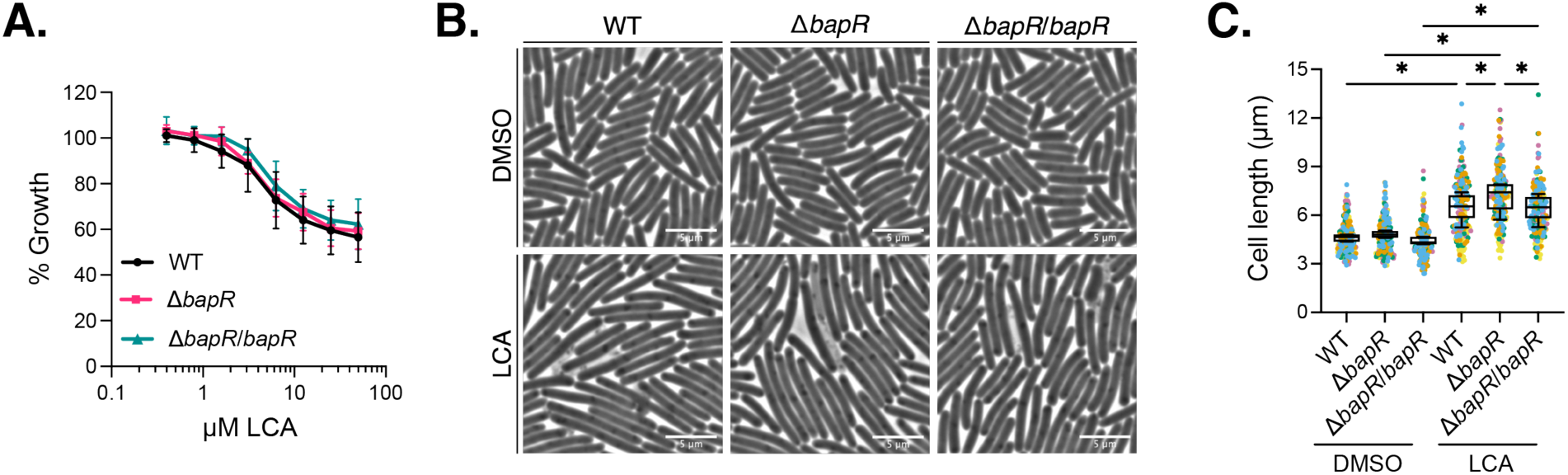
BapR influences cell length in the presence of LCA. (**A**) Inhibition of WT, Δ*bapR*, and Δ*bapR*/*bapR C. difficile* growth in broth culture across various concentrations of LCA calculated from OD_600_ after 5 hours of growth; error bars represent SD, n = 2. (**B**) Phase-contrast images of wildtype (WT) *C. difficile*, Δ*bapR*, and Δ*bapR*/*bapR,* the complement strain carrying *bapR* at an ectopic locus after 3-hour treatment with 20 μM LCA or DMSO vehicle during log-phase; images are representative of 5 independent experiments. (**C**) Measurement of cell length from the experiments in (A); at least 460 cells were measured per strain per condition for each experiment and the length of 35 random cells per experiment are shown (75). Colors represent independent experiments, *p < 0.05 by repeated measures one-way ANOVA with Tukey correction.

To gain initial mechanistic insight into BapR’s response to bile acids in *C. difficile*, we asked whether *bapR* expression changes during bile acid-mediated stress. To this end, we measured *bapR* transcript levels by qRT-PCR after 1-hour exposure to 20 μM LCA or DMSO vehicle during log-phase. Expression of *bapR* decreased ∼3-fold in the WT strain upon LCA treatment (*bapR* transcript was undetectable in the Δ*bapR* strain as expected) (**Figure 4A**). Despite the unanticipated overexpression of *bapR* in the Δ*bapR*/*bapR* complement strain, *bapR* expression was down-regulated upon treatment of these cells with LCA (**Figure 4A**).

**Figure 4.**
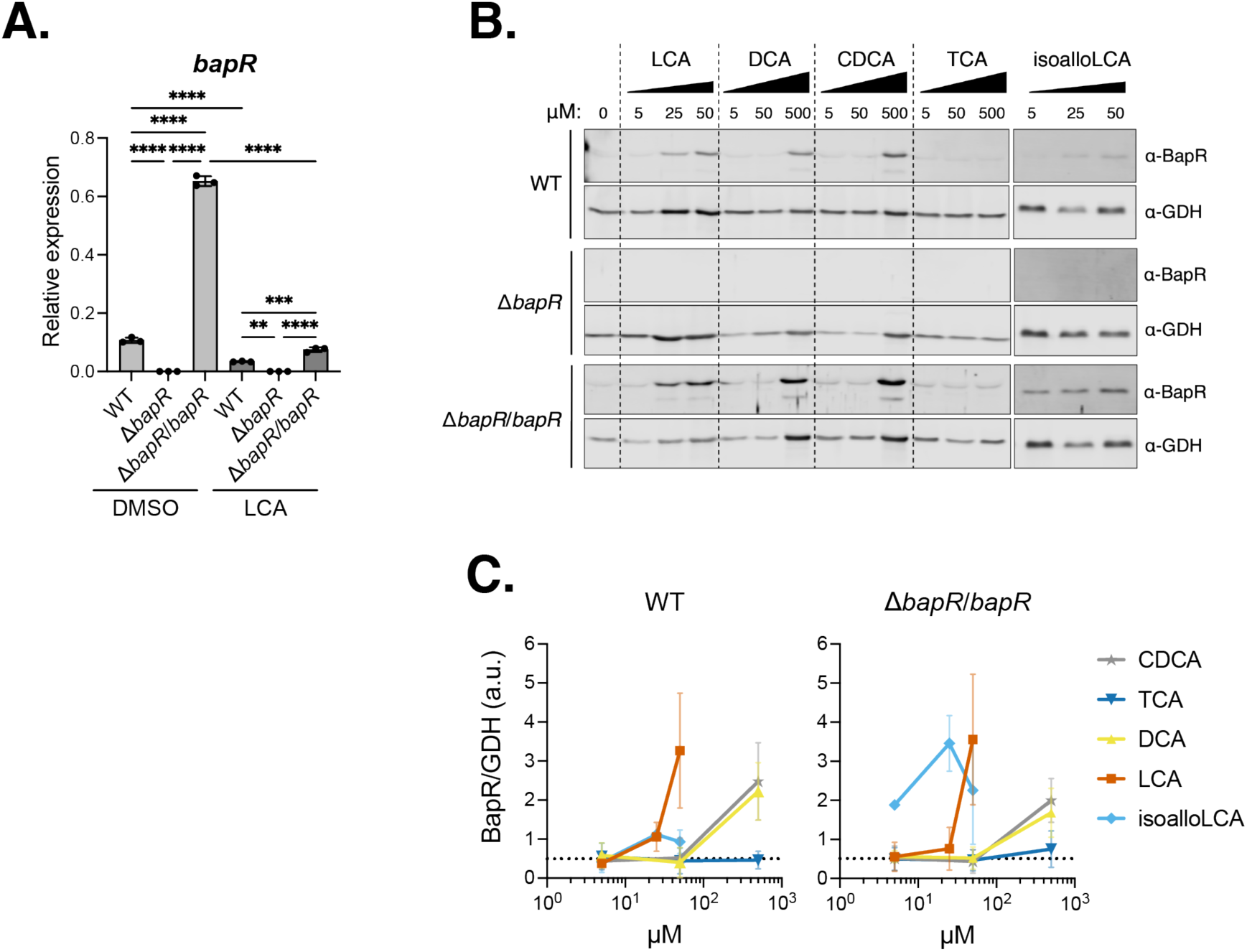
BapR is stabilized by bile acids. (**A**) Expression of *bapR* measured by qRT-PCR after 1-hour exposure to 20 μM LCA or DMSO vehicle; expression is relative to the housekeeping threonyl-tRNA synthetase *thrS* (76). n = 3, **p < 0.01, ***p < 0.001, ****p < 0.0001 by one-way ANOVA with Tukey correction. (**B**) Bile acids were added to log-phase *C. difficile* cultures at the indicated concentrations and samples were taken after 3 hours for Western blotting; glutamate dehydrogenase (GDH) serves as a loading control and blots are representative of 3 biological replicates. (**C**) Quantification of the blots in (A); n = 3.

Since our finding that *bapR* expression decreased in response to LCA was somewhat surprising, we wondered whether this decrease corresponded to lower protein levels. To test this possibility, we treated log-phase *C. difficile* cultures with increasing concentrations of LCA, DCA, CDCA, TCA, or isoalloLCA for 3 hours during log-phase and assessed BapR levels by Western blot. Unexpectedly, we observed a dose-dependent increase in BapR levels with LCA, and to a lesser extent with DCA, CDCA, and isoalloLCA (**Figure 4B and C**). This implies that BapR is stabilized by these bile acids or becomes less susceptible to degradation in *C. difficile*, since *bapR* transcript levels are decreased upon LCA treatment (**Figure 4A**). The elevated BapR levels were sustained for at least 6 hours of growth with LCA (**Figure S4**). In agreement with the apparent binding affinity measured in thermal shift assays, ∼10-fold higher concentrations of DCA and CDCA were needed to stabilize BapR at levels comparable to LCA (∼3-fold increase at 50 μM LCA vs. ∼2-fold increase at 500 μM DCA or CDCA). TCA did not change BapR levels even at 500 μM, and since BapR did not appreciably bind TCA in our thermal shift assays, the elevated BapR levels in *C. difficile* seen upon treatment with the other bile acids is likely due to binding. It is unclear why isoalloLCA increased BapR levels less than LCA in WT *C. difficile* and more in the complementation strain, but these data nevertheless suggest that BapR interacts with bile acids in *C. difficile*.

### BapR controls gene expression, in some cases in an LCA-dependent manner

Since BapR is predicted to be a MerR-type transcription factor, we next asked whether it regulates gene expression, particularly in an LCA-dependent manner. To this end, we assessed the transcriptome of WT and Δ*bapR C. difficile* during short-term LCA exposure using RNA-seq. Log-phase cultures of each strain were treated with 20 μM LCA or DMSO vehicle for 1 hour before harvesting RNA for next-generation sequencing. Transcriptomic analysis revealed that LCA induced global changes in gene expression (569 genes significantly upregulated and 580 genes significantly downregulated; p < 0.05 and fold change > 2) (**Figure S5A** and **Table S2**). We noted changes in line with previously reported effects of LCA on *C. difficile*: motility genes were largely downregulated in our analyses, consistent with LCA causing loss of flagella (38) (**Figure S5B**). While the expression of many genes was altered by LCA, we were most interested in genes whose expression was modulated in a BapR-dependent manner. We found that BapR significantly influenced the expression of three gene loci: *mdeA-cd3576* (CD630_35770-CD630_35760), *cd0618*-*cd0616* (CD630_06180-CD630_06160), and *cwpV* (CD630_05140) (**Figure 5A**). Expression of these genes was higher in the Δ*bapR* strain relative to WT (2-4-fold different), indicating that BapR represses them.

**Figure 5.**
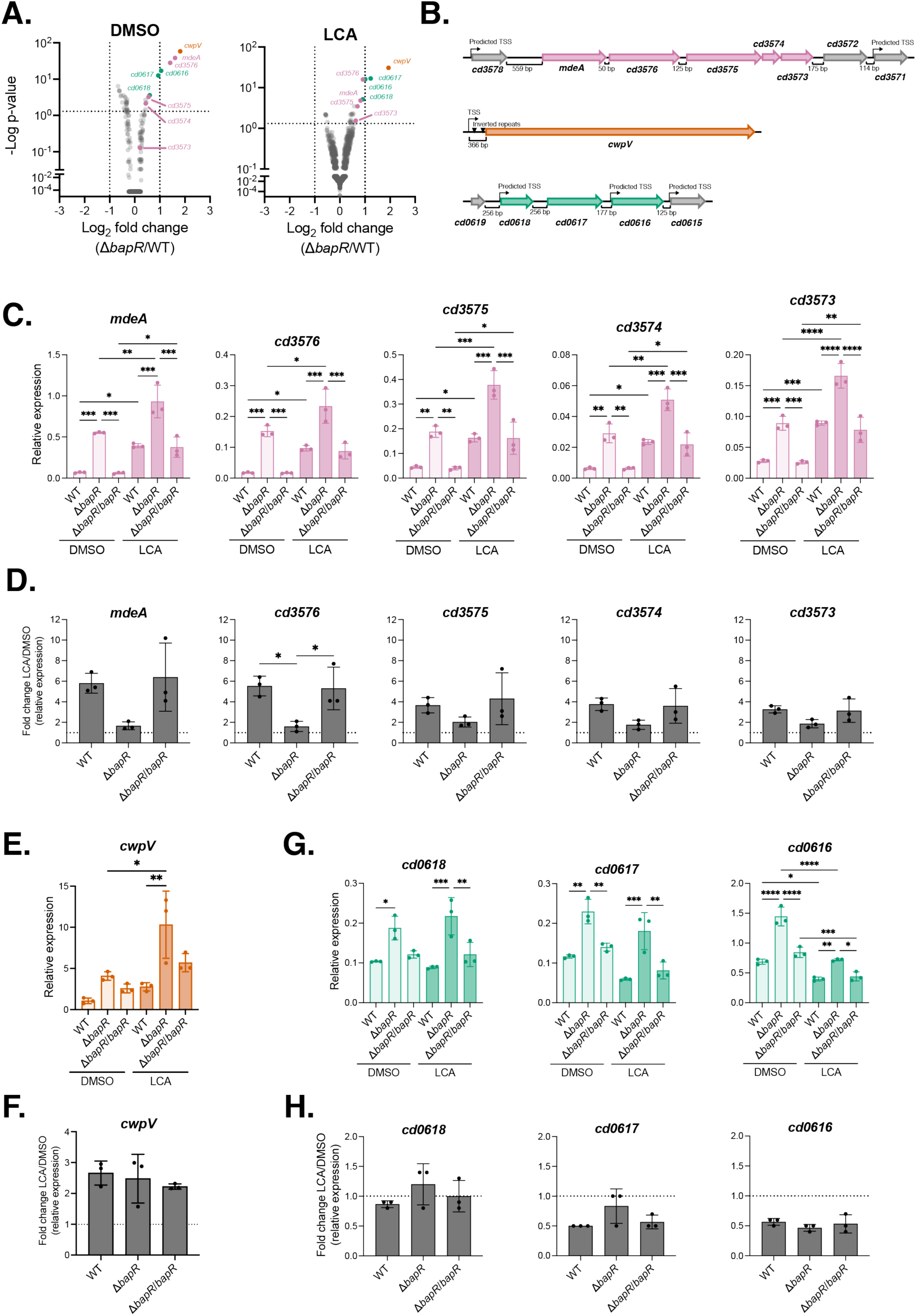
Genes regulated by BapR. (**A**) RNA-seq analysis of WT and Δ*bapR C. difficile* after 1-hour treatment with DMSO vehicle or 20 μM LCA during log-phase; dashed lines indicate significance cutoffs at p < 0.05 and fold change>2, n = 3. (**B**) Genomic context of hits from (A). (**C**) Expression of the *mdeA* gene cluster measured by qRT-PCR using purified RNA that is distinct from the 1-hour exposure to DMSO or 20 μM LCA used for RNA-seq; expression is relative to *thrS*, n = 3. (**D**) Relative expression data in (C) plotted as fold change LCA over DMSO for each strain. (**E**) Expression of *cwpV* as in (C). (**F**) Fold change LCA/DMSO for *cwpV* as in (D). (**G**) Expression of *cd0618* gene cluster as in (C). (**H**) Fold change LCA/DMSO for *cd0618* gene cluster as in (D). *p < 0.05, **p < 0.01, ***p < 0.001, ****p < 0.0001 by one-way ANOVA with Tukey correction.

Of the genes significantly affected by BapR, *mdeA* and *cd3576* expression changed the most between DMSO and LCA conditions. These genes were differentially expressed in the DMSO control (WT vs. Δ*bapR*) but not in the presence of LCA (**Figure 5A**). *mdeA* encodes a methionine γ-lyase thought to be responsible for methanethiol and H_2_S production from methionine and homocysteine/cysteine, respectively, (55). *cd3576* encodes a major facilitator superfamily (MFS) transporter. These genes are found in close proximity to the upstream genes, *cd3575-cd3573* (**Figure 5B**), which are predicted to encode two hypothetical proteins and a sodium:solute symporter, respectively. While genome-wide transcription start site (TSS) mapping in *C. difficile* predicts promoters at *cd3578* and *cd3571*, no TSSs were reported for *mdeA*-*cd3572* (56) (**Figure 5B**), although in our experience these genome-wide analyses do not capture all TSS. Since *cd3573*-*cd3575* approached significance under at least one condition (*cd3572* and *cd3578* did not) and may be co-transcribed with *cd3576* and *mdeA* based on proximity and orientation, we validated expression of *mdeA-cd3573* using qRT-PCR. In line with our RNA-seq analyses, we found that all genes in this cluster were expressed more in the Δ*bapR* strain than WT under both conditions, and the differential expression was complemented in the Δ*bapR*/*bapR* strain (**Figure 5C**). Additionally, LCA increased the expression of all genes in the cluster across all strains (**Figure 5C**). To compare the response to LCA between strains, we plotted expression levels for a given strain as the fold-change in LCA relative to DMSO (LCA/DMSO). This revealed that the loss of BapR resulted in a smaller induction of these genes in the presence of LCA relative to WT levels (**Figure 5D**); the differential response was most apparent for *mdeA* and *cd3576* (∼6-fold induction in the WT and complement strains vs. ∼1.5-fold induction in the Δ*bapR* strain). Taken together these data suggest that BapR is necessary to induce these genes in response to LCA, likely by de-repressing their expression upon sensing LCA.

In contrast, *cwpV*, which was differentially expressed between Δ*bapR* and WT *C. difficile* in our RNA-seq analyses, was not affected by LCA treatment (**Figure 5A**). CwpV is a cell wall protein that makes up ∼13% of the total surface layer proteins in *C. difficile* (57). It promotes aggregation of *C. difficile* cells *in vitro* (57) and confers resistance to *Siphoviridae* and *Myoviridae* family phages (58), but is expressed by only ∼5% of cells in culture due to a phase-variable RecV-controlled genetic switch located between its promoter and coding DNA sequence (59). In bulk population measurements by qRT-PCR, *cwpV* expression was higher in the Δ*bapR* strain than WT and the complement, particularly in the presence of LCA (**Figure 5E**). However, the fold-change in *cwpV* expression induced by LCA (∼2.5-fold in WT) was the same for Δ*bapR* and the Δ*bapR*/*bapR* complementation strain (**Figure 5F**). These data suggest that BapR indirectly represses *cwpV*, since the LCA-induced upregulation of *cwpV* is not dependent on BapR. Given that LCA causes global changes in the *C. difficile* transcriptional landscape (**Figure S5**), other unknown factors likely mediate the LCA-induced expression of *cwpV*.

Our RNA-seq analyses also identified a second cluster of genes whose expression changed in a BapR-dependent manner: *cd0618-0616*. The differential expression of *cd0616* and *cd0617* hovered at the significance cutoff, and *cd0618* approached significance under both conditions (**Figure 5A**). *cd0618* encodes a LytTR family transcription factor; *cd0617* encodes a CPBP family intramembrane metalloprotease; and *cd0616* encodes another MerR family transcription factor. TSSs were previously predicted for *cd0616* and *cd0618*, but not for *cd0617* (56) (**Figure 5B**). qRT-PCR analyses confirmed that these genes are over-expressed in the Δ*bapR* strain relative to WT and the complement strain (**Figure 5G**), but only the expression of *cd0616* and *cd0617* was affected by LCA treatment (∼2-fold reduction) (**Figure 5H**). LCA had similar effects on the expression of these genes between strains irrespective of whether BapR was present, indicating that like *cwpV,* BapR does not control their LCA-dependent expression (**Figure 5H**). While it is unclear whether there are other conditions in which BapR derepresses the expression of *cwpV* or the *cd0618-cd0616* locus, our data nonetheless imply that (i) BapR represses gene expression directly and indirectly and that (ii) separate, unknown mechanisms of LCA-dependent transcriptional regulation act in parallel to BapR.

### BapR binds the promoter region of *mdeA* but not *cd0616*

To test whether BapR directly regulates the *mdeA* cluster, *cd0618* cluster, or *cwpV*, we performed electrophoretic mobility shift assays (EMSAs) with the promoter regions of these genes. These DNA fragments were fluorescently labeled at their 5’ ends and incubated with purified BapR. Dose-dependent mobility shifts of the *mdeA* promoter region were observed for BapR, and importantly, this shift was competed away by an excess of the same promoter region lacking the fluorescent label (cold competitor; **Figure 6A**). In line with our transcriptional analyses suggesting that BapR indirectly regulates *cwpV*, no shift was seen for the *cwpV* promoter in its “on” orientation (**Figure 6B**) Further, BapR did not bind the promoters of *cd0618* or *cd0616* (**Figure S6**). BapR failed to bind its own promoter region indicating that it does not autoregulate (**Figure S6**).

**Figure 6.**
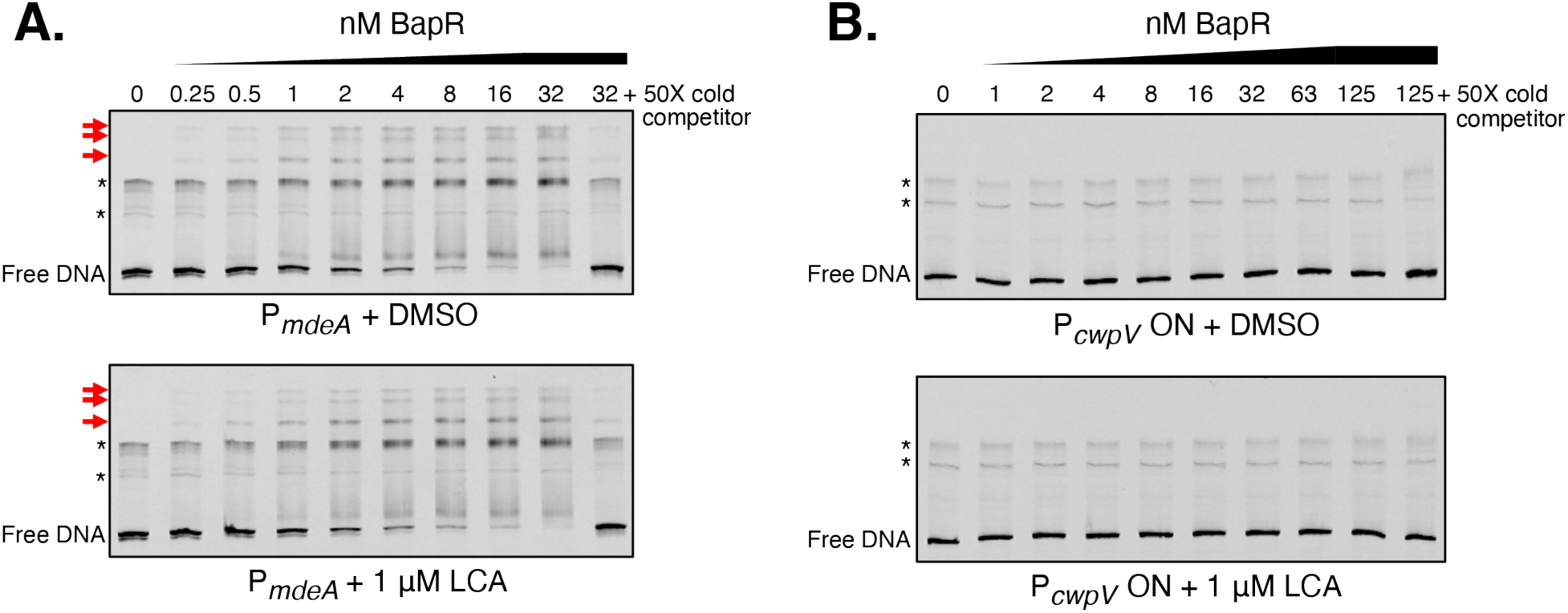
BapR directly regulates *mdeA*. (**A**) Electrophoretic mobility shift assay with purified BapR and a 250 bp DNA fragment comprising the region immediately upstream of the *mdeA* start codon as a putative promoter; 20 fmol 5’ IRDye800-labeled promoter fragment per lane, the last lane contains 20 fmol labeled DNA and 1,000 fmol of the same DNA fragment lacking the fluorescent label as a cold competitor. Gel is representative of 3 replicates with protein from 2 independent purifications. “Free DNA” indicates unbound DNA and red arrows indicate BapR-bound DNA. Asterisks denote nonspecific bands that likely represent different DNA secondary structures. (**B**) Assay as in (A) with a 356 bp DNA fragment encompassing the *cwpV* promoter in its “on” orientation.

Notably, we found that LCA addition to the EMSAs did not affect BapR binding to the *mdeA* promoter (**Figure 6A**). This result was not necessarily surprising given that MerR family transcription factors remain bound to their DNA targets regardless of whether their C-terminal domain has bound their ligands. Instead, ligand binding results in these transcription factors regulating transcription via DNA distortion to reorient the -35 and -10 sites and facilitate productive RNA polymerase interactions (51, 60, 61). Since MerR family proteins can act as repressors in the absence of ligand and activators in the presence of ligand, our finding that BapR binds the *mdeA* promoter independently of LCA (**Figure 6A**) is consistent with previously characterized members of this family (60, 62–64). Furthermore, since BapR maintains the ability to bind DNA even in the presence of a large molar excess of LCA (**Figure 6A**), the reduced thermal stability of BapR observed in the presence of LCA (**Figure 2D**) is consistent with BapR undergoing a conformational change, rather than being destabilized by LCA.

## Discussion

In this study, we identified BapR as a novel sensor for bile acids in *C. difficile* that de-represses the expression of a small subset of genes upon sensing LCA-related bile acids. Our chemical proteomics, affinity pull-downs, and thermal shift assays indicate that BapR specifically binds bile acids, especially LCA (**Figure 2**). Our data further suggest that BapR is stabilized by bile acids in *C. difficile* (**Figure 4**), which allows in BapR to directly de-repress the expression of genes encoding the methionine γ-lysase MdeA and two putative transporters. Specifically, our data indicate that BapR binds the promoter region of *mdeA* and represses the expression of the *mdeA* gene cluster in the absence of LCA; however, upon binding LCA, BapR undergoes a conformational change that reorients the *mdeA* promoter and licenses transcription (**Figure 7**). Given MdeA’s involvement in sulfur metabolism (55) and the fact that bile acids are found exclusively in the gut of metazoans, we hypothesize that this LCA-sensing system regulates *C. difficile*’s metabolic adaptation to the gut environment. Since cysteine is elevated ∼6-fold in the dysbiotic murine cecum (5), it is plausible that *C. difficile* would upregulate cysteine catabolism in this environment. Other factors downstream of this metabolic adaptation may be responsible for the cell length phenotype (**Figure 3**), but further studies are needed to delineate connections between these observations. Our RNA-Seq and qRT-PCR analyses also revealed that the expression of genes encoding the cell surface protein, CwpV, two other transcription factors, and a putative metalloprotease is repressed by BapR (**Figures 5 and 6**), but conditions under which BapR may de-repress these genes remain to be determined.

**Figure 7.**
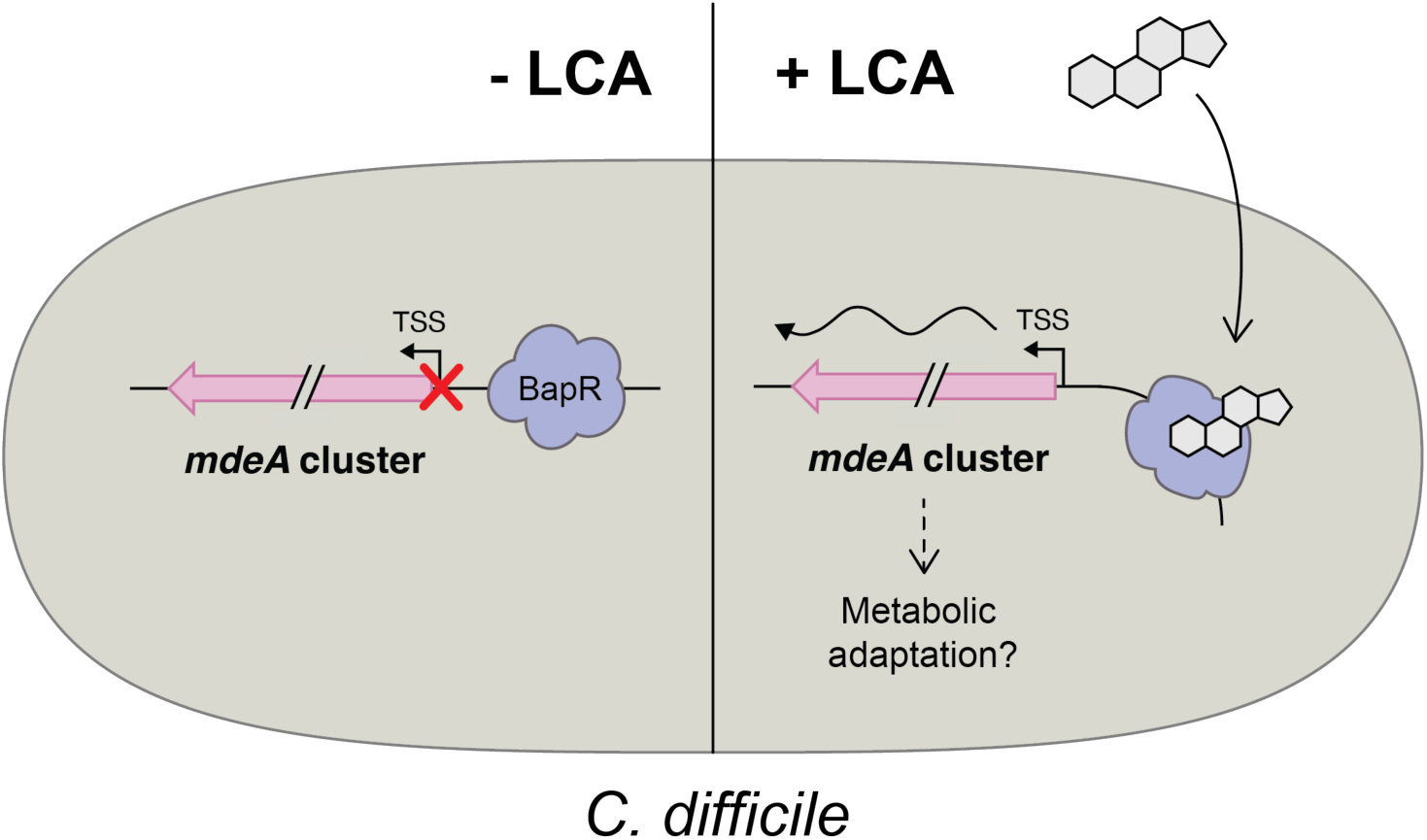
Model of proposed BapR function. Consistent with MerR family proteins, BapR is DNA-bound in the absence of LCA and represses gene expression. Upon binding LCA, BapR changes conformation to reorient the promoter and allow transcription, possibly for the purpose of metabolic adaptation.

Although BapR does not appear to be involved in resistance to LCA in the broth culture-based growth conditions tested, it is possible that the fitness of the Δ*bapR* strain would be reduced in competition with WT *in vitro* or *in vivo*. It is also possible that *C. difficile* has factors redundant to BapR that compensate for its loss. However, the strong correlation between presence of DCA and LCA in the gut and resistance to CDI could indicate that *C. difficile* does not have effective resistance mechanisms for these microbiota-generated metabolites. Instead, *C. difficile* appears to grow opportunistically in gut environments with very low levels of DCA and LCA (4–7), where these low levels could serve as environmental cues. Another possibility is that other stressors in the gut milieu could potentiate LCA’s toxicity and reveal a resistance role for BapR. For example, antibiotics produced by 7α-dehydroxylating bacteria like *C. scindens*, which secretes 1-acetyl-β-carboline, and *Paraclostridium sordellii*, which produces turbomycin A and 1,1,1-tris(3-indolyl)-methane (TIM), are all active against *C. difficile* (36) and could potentiate the toxicity of LCA. While we attempted to test this hypothesis using the commercially available 1-acetyl-β-carboline, we unfortunately found that *C. difficile* strain 630Δ*erm* is substantially more resistant than the ATCC 9689 strain used in the study described above (36) (such that solubility limited our ability to test our hypothesis). Regardless, investigating the interactions between LCA-producing bacteria and *C. difficile* could reveal that LCA sensing by BapR gives rise to adaptations that confer a competitive advantage for *C. difficile*.

Determining the functions of the other genes repressed by BapR in an LCA-dependent manner may provide further insight into the function of this MerR family member. Notably, most MerR family transcription factors sense toxic molecules and upregulate cognate efflux pumps accordingly. Since *cd3575* is annotated as a sodium:solute symporter and *cd3576* a MFS transporter, a plausible function of this system is to transport metabolites related to cysteine catabolism. However, most bile acid-binding transcription factors in bacteria regulate production of multidrug efflux pumps (65–68), and CD3575 and CD3576 could play a role in bile acid efflux. To date a transporter or channel for bile acids has not been identified in *C. difficile*, but we can infer this phenomenon from our chemical proteomics screen as the probes were added to intact cells in culture, washed away, then UV crosslinked. Indeed, many of the hits from our screen are cytosolic proteins (**Table S1**). A few examples of bacterial bile acid transporters have been described: the *baiG* gene found in 7α-dehydroxylating *Clostridium* and *Eubacterium* species encodes a bile acid transporter (29, 69), *Lactobacillus johnsonii* possesses a bile acid MFS transporter in conjunction with bile salt hydrolase activity (70), and *Neisseria meningitidis* and *Yersinia frederiksenii* both have a homologue of the eukaryotic ASBT sodium:bile acid co-transporter that imports TCA (71–74). *C. difficile* CD3575 and CD3576 share only 11-14% identity with each of these bile acid transporters, but the possibility remains that they export bile acids.

Only a few other bacterial transcription factors have been shown to directly sense bile acids (65–68), and to our knowledge BapR is the only example that detects bile acids for metabolic adaptation. While our data are consistent with BapR being a bile acid sensor, *C. difficile* almost certainly encodes alternative mechanisms for sensing or responding to LCA. Global transcriptional changes occur in the presence of LCA (**Figure S5**), yet BapR regulates the expression of only a handful of genes (**Figure 5**). Indeed, our chemical proteomics screen detected a few other putative transcription factors, two-component system histidine kinases, and serine/threonine kinases as candidate bile acid-binding proteins with potential signaling roles. Future studies of these hits could reveal pathways by which *C. difficile* deals with bile acid stress independent of BapR. For example, while we did not biochemically validate essential hits from our proteomics screen, inactivation of these proteins by LCA could explain its toxicity. Thus, it remains to be seen which are involved and whether the toxicity is due to the concerted inactivation of multiple proteins. Additional studies using bile acid photoaffinity probes and chemical proteomics should reveal other protein targets and mechanisms of action for these prominent gut microbiota metabolites.

## Materials and Methods

### Bacterial strains and growth conditions

*C. difficile* strains are of the 630Δ*erm* background and mutants were constructed using *pyrE*-based allele-coupled exchange (54). Strains were grown on brain heart infusion medium supplemented with 0.5% w/v yeast extract and 0.1% w/v L-cysteine (BHIS) with taurocholate (TCA; 0.1% w/v; 1.9 mM), thiamphenicol (10–15 μg/mL), kanamycin (50 μg/mL), or cefoxitin (8 μg/mL) as needed. *C*. *difficile* defined medium (CDDM) (77)was supplemented with 5-fluoroorotic acid at 2 mg/mL and uracil at 5 μg/mL. Cultures were grown swirling at 37°C under anaerobic conditions using a gas mixture containing 85% N_2_, 5% CO_2_ and 10% H_2_. *Escherichia coli* strains were grown at 37°C with shaking at 225 rpm in Luria-Bertani medium (LB) or at 20°C with shaking at 225 rpm in autoinduction broth (Terrific broth [Thermo Fisher] with 0.5% glycerol, 0.05% glucose, and 0.1% α-lactose monohydrate). Media were supplemented with chloramphenicol (20 μg/mL), ampicillin (100 μg/mL), or kanamycin (30 μg/mL) as needed.

### *C. difficile* growth curves

*C. difficile* cultures were grown for 3 hours, back-diluted 1:50, and grown for an additional 3 hours to an OD_600_ of ∼0.3. Cultures were diluted to an OD_600_ of 0.05 and added to 96-well plates with 2X pre-reduced bile acids or probes in BHIS for a total volume of 150 μL/well (75 μL culture + 75 μL 2X compound in BHIS). OD_600_ was read every 10 minutes for 18 hours with constant shaking at 37°C in an Epoch 2 plate reader (BioTek) in an anaerobic chamber. Percent growth inhibition was calculated from the OD_600_ relative to DMSO at 5 hours. Two biological replicates from independent starter cultures were used per experiment.

### In-gel fluorescent bile acid probe labeling

*C. difficile* cultures were grown for 3 hours, back-diluted 1:25 into 25 mL BHIS, and grown for an additional 3-4 hours to an OD_600_ of ∼0.7. Bile acid probes or DMSO were added to the cultures and incubated for 30 minutes. Cells were resuspended in 1 mL PBS, transferred to a 24-well plate, and UV irradiated (365 nm) uncovered for 5 minutes on ice 3-5 cm from the lamp. Probe-crosslinked cells were pelleted, resuspended in 500 μL cold lysis buffer (1X Halt protease inhibitor [Thermo Fisher], 0.5 mg/mL lysozyme, 1:1000 benzonase [EMD Millipore], and 0.1% NP-40 in 1X PBS). Cells were lysed by four 30-second rounds of bead beating at speed 6 with Lysing Matrix B (MP Bio) in a Fast Prep-24 bead beater (MP Bio). Beads were pelleted at 3,000g for 2 minutes and the total lysate was collected. Alexa Fluor 488 was conjugated to the probe in 30 μg total lysate using a Click-iT Plus Alexa Fluor 488 Picolyl Azide Toolkit (Thermo Fisher). Protein was precipitated overnight at -20°C in methanol, washed twice with cold methanol, and resuspended in 25 μL SDS-PAGE sample buffer. Fluorescent probe labeling was visualized in-gel following SDS-PAGE using the SYBR Safe long pass blue filter in a Fujifilm FLA-9000 imager at 200 μM resolution. The gel was Coomassie stained as a loading control.

### Chemical proteomics

*C. difficile* overnight cultures were diluted 1:25 into 20 mL BHIS and grown to an OD_600_ of 1.1. Bile acid probes or DMSO were added at 10 μM for 1 hour. Cells were resuspended in 1 mL phosphate-buffered solution (PBS), transferred to a 24-well plate, and UV irradiated (365 nm) uncovered for 5 minutes on ice 3-5 cm from the lamp. Probe-crosslinked cells were pelleted, resuspended in 500 μL cold lysis buffer (1X Halt protease inhibitor [Thermo Fisher], 0.5 mg/mL lysozyme, 1:1000 benzonase [EMD Millipore], and 0.1% NP-40 in 1X PBS). Cells were lysed by four 30-second rounds of bead beating at speed 6 with Lysing Matrix B (MP Bio) in a Fast Prep-24 bead beater (MP Bio). Beads were pelleted at 3,000 g for 2 minutes and the total lysate was collected. The lysate was flash-frozen in liquid nitrogen before further processing. Cell lysates were centrifuged at 16000 g for 20 min to remove cell debris and supernatants were collected. Each total cell lysates was added with 100 μL of click chemistry reagents as a 10X master mix (az-Biotin: 0.1 mM, 10 mM stock solution in DMSO; tris(2-carboxyethyl)phosphine hydrochloride (TCEP): 1 mM, 50 mM freshly prepared stock solution in dH_2_O; tris[(1-benzyl-1H-1,2,3-triazol-4-yl)methyl]amine (TBTA): (0.1 mM, 2 mM stock in 4:1 *t*-butanol: DMSO); CuSO_4_ (1 mM, 50 mM freshly prepared stock in dH_2_O). Samples were mixed well and incubated at room temperature for 1 h. After incubation, samples were mixed with 4 mL cold methanol and incubated at -20 °C overnight. Protein pellets were centrifuged at 5000 g for 30 min at 4°C, pellets were transferred to 2.0 mL centrifuge tube and were washed with 1 mL cold methanol 3 times. After last wash, pellets were let air dried before being re-solubilized in 250 μL 4% SDS PBS with bath sonication. Solutions were diluted with 750 μL PBS, and incubated with 100 μL PBS-T-washed High Capacity NeutrAvidin agarose (Pierce) (500 μL PBS-T-washed twice, 2500 g for 60 s) at room temperature for 1 h with end-to-end rotation. The agarose was washed with 500 μL 1% SDS PBS 3 times, 500 μL 1M Urea PBS 3 times, and 500 μL PBS, 3 times and then reduced with 500 μL 10 mM DTT (Sigma) in PBS for 30 min at 37 °C, and alkylated with 500 μL 50 mM iodoacetamide (Sigma) in PBS for 30 min in dark. 50 μL NH_4_HCO_3_ (10 mM) was added to the tube. Neutravidin-bound proteins were digested on bead with 400 ng Trypsin/Lys-C mix (Promega) at 37 °C overnight with shaking. Digested peptides were collected (2500 g for 60 s) and lyophilized before being desalted with custom-made stage-tip containing Empore SPE Extraction Disk (3M). Peptides were eluted with 2% acetonitrile, 2% formic acid in dH_2_O.

Peptide LC-MS analysis was performed with a Dionex 3000 nano-HPLC coupled to an Orbitrap XL mass spectrometer (Thermo Fisher). Peptide samples were pressure-loaded onto a home-made C18 reverse-phase column (75 µm diameter, 15 cm length). A 180-minute gradient increasing from 95% buffer A (HPLC grade water with 0.1% formic acid) and 5% buffer B (HPLC grade acetonitrile with 0.1% formic acid) to 75% buffer B in 133 minutes was used at 200 nL/min. The Orbitrap XL was operated in top-8-CID-mode with MS spectra measured at a resolution of 60,000@m/z 400. One full MS scan (300–2000 MW) was followed by three data-dependent scans of the most intense ions with dynamic exclusion enabled. Label-free quantification of bile acid probe-labeled proteins was performed in MaxQuant software as described (78). The search results from MaxQuant were analyzed by Perseus (http://www.perseusframework.org/). Briefly, the DMSO and bile acid probe-labeled replicates were grouped correspondingly. The results were cleaned to filter off reverse hits and contaminants. Only proteins that were identified in 3 out of 4 sample replicates and with more than two unique peptides were subjected to subsequent statistical analysis. LFQ intensities were used for measuring protein abundance and logarithmized (base 2). Signals that were originally zero were imputed with random numbers from a normal distribution, whose mean and standard deviation were chosen to best simulate low abundance values below the noise level (Normal distribution: Width = 0.3; Shift = 1.8).

### Bile acid probe pull-downs of BapR

*C. difficile* cultures were grown for 4 hours, back-diluted 1:2,000 into 70 mL BHIS, and grown overnight. Cultures were then diluted to an OD_600_ of 1, split into 20 mL/condition and incubated with probe or DMSO for 1 hour. Cells were resuspended in 1 mL PBS, transferred to a 6-well plate, and UV irradiated (365 nm) uncovered for 5 minutes on ice 3-5 cm from the lamp. Probe-crosslinked cells were pelleted, resuspended in 1 mL cold lysis buffer (1X Halt protease inhibitor [Thermo Fisher], 0.5 mg/mL lysozyme, 1:1000 benzonase [EMD Millipore], and 0.1% NP-40 in 1X PBS). Cells were lysed by four 30-second rounds of bead beating at speed 6 with Lysing Matrix B (MP Bio) in a Fast Prep-24 bead beater (MP Bio). Beads were pelleted at 21,000g for 5 minutes at 4°C and the cleared lysate was collected. Biotin was conjugated to the probes in a click reaction with 0.4 mg cleared lysate in 180 μL PBS and 20 μL 10X click master mix (1 mM azido-PEG3-biotin [Alfa Aesar], 10 mM tris(2-carboxyethyl)phosphine hydrochloride, 1 mM tris[(1-benzyl-1H-1,2,3-triazol-4-yl)methyl]amine in 4:1 t-butanol:DMSO, and 10 mM copper sulfate pentahydrate) and protein was precipitated overnight at -20°C in methanol. Precipitates were washed twice with cold methanol, dried at 37°C for 1 hour, and resolubilized in 50 μL 4% sodium dodecyl sulfate (SDS) in PBS with bath sonication. 150 μL PBS was added and a sample was taken as input before incubation with PBS+0.1% Tween-20-washed Pierce High Capacity NeutrAvidin agarose beads (Thermo Fisher) for 1 hour at room temperature with end-over-end rotation. Beads were washed three times each with PBS+1% SDS, PBS+4M urea, then PBS and boiled to elute biotin-probe-protein complexes.

### Recombinant BapR purification

BL21(DE3) *E. coli* encoding lactose-inducible *bapR* with a C-terminal autoprocessing CPD-His tag was grown in 20 mL LB with ampicillin, then back-diluted 1:1,000 into 1L autoinduction broth with ampicillin and grown at 20°C for ∼60 hours. Cultures were pelleted, resuspended in 50 mL low imidazole buffer (LIB; 500 mM NaCl, 50 mM Tris-HCl pH 7.5, 15 mM imidazole, 10% glycerol, 2 mM β-mercaptoethanol), and flash frozen in liquid nitrogen. Once thawed, cells were probe sonicated (Branson) in 3 x 45 second rounds at 40% amplitude with 5 minutes on ice between. Lysates were cleared by centrifugation at 13,000 rpm for 45 min at 4°C. BapR-CPD-His was affinity purified from cleared lysates using Ni-NTA agarose beads (Qiagen) with gentle rocking at 4°C for 2 hours. The beads were washed three times with LIB before inducing cleavage of the CPD tag (79) with 200 μM inositol hexakisphosphate in LIB at 4°C overnight to elute untagged BapR. The eluted protein was buffer-exchanged into SEC buffer (200 mM NaCl, 10 mM Tris-HCl pH 7.5, 5% glycerol, and 1 mM dithiothreitol) and concentrated using an Amicon Ultra-15 10 kDa cutoff centrifugal filter (Millipore Sigma). Affinity-purifued protein was further purified by size exclusion chromatography (SEC) using a Superdex 200 Increase 10/300 GL column (GE) on an AKTA Pure fast protein liquid chromatography instrument (GE), reconcentrated, and flash frozen in aliquots.

### Thermal shift assays

Affinity- and SEC-purified BapR was added to a mix of 5X SYPRO Orange dye (Thermo Fisher) and the indicated concentrations of bile acids or DMSO in 1.5X PBS to a final concentration of 1 μM in a 96-well white bottom plate. Fluorescence was measured as temperature increased 1°C/minute from 25°C to 95°C in a StepOne Plus qPCR instrument (Applied Biosystems). Protein from 2 independent purifications was used, and a no-protein control for each ligand at each concentration was used to identify cutoffs above which the ligand generated background fluorescence, if at all.

### Phase-contrast microscopy

*C. difficile* was inoculated into BHIS cultures and grown for 3 hours, back-diluted 1:50, and grown for another 3 hours. Cultures were then split and treated with DMSO or bile acids at the indicated concentrations for 3 hours. Aliquots of the cultures were pelleted, resuspended in ∼20 μL PBS, and 1 μL was spotted on a 1% agarose pad poured in a Gene Frame (Thermo Fisher). Pads were sealed with a coverslip inside the anaerobic chamber. Phase images were acquired on a Zeiss Axioskop using a 100X Plan-NEOFLUAR oil phase objective. Cell length was measured from pole-to-pole using Fiji software and at least 460 cells were measured per strain/condition in 5 independent experiments.

### BapR protein induction by bile acids in *C. difficile*

*C. difficile* starter cultures were grown for 3 hours, back-diluted 1:50 into 30 mL BHIS, and grown for an additional 3 hours before being split into new tubes with the indicated concentrations of bile acids or DMSO. After 3 hours cells were pelleted, resuspended in sample loading buffer, frozen at -20C, and boiled before running SDS-PAGE. Protein was transferred to a PVDF membrane, blocked with 0.5X blocking buffer (LI-COR), and probed with chicken α-GDH antibody (Thermo Fisher) and a custom mouse α-BapR antibody (Cocalico Biologicals). Antibodies were detected using IRDye700- or IRDye800-conjugated donkey α-chicken and goat α-mouse secondary antibodies (LI-COR). The blots were imaged using a LI-COR Odyssey CLx imager and quantified using ImageStudio Lite software (LI-COR).

### RNA extraction, RNA-seq, and qRT-PCR

*C. difficile* cultures were grown for 3-4 hours, back-diluted 1:50, and grown for an additional 3-4 hours before being split for treatment with DMSO or 20 μM LCA. After 1 or 3 hours of LCA exposure (OD_600_ of 0.1-0.2 at 1 hour or 0.2-0.5 at 3 hours) RNA was extracted using a FastRNA Pro Blue kit (MP Bio). Samples were treated twice for 45 minutes at 37°C with DNase I (New England Biolabs) with heat inactivation at 75°C for 10 minutes and DNA was further removed using a RNeasy Mini kit (Qiagen). RNA was harvested from three independent cultures as biological replicates per experiment, and independent extractions were used for RNA-seq and qRT-PCR validation.

For RNA-seq analysis an Agilent Bioanalyzer was used to verify RNA quality before depleting rRNA and ligating adapters and indexes using a Stranded Total RNA with RiboZero Plus library preparation kit (Illumina). Samples were sequenced as single-end 75 reads on an Illumina NextSeq 500 sequencer at the Tufts University Genomics Core Facility. Sequences were trimmed using BBDuk, mapped to the *C. difficile* 630 genome, and analyzed for differential expression using DESeq2 in Geneious Prime software. Gene functional characterization was done using GSEA-Pro v3 (University of Groningen) to classify by COG terms and manual classification for genes that were not classified by GSEA-Pro.

For qRT-PCR analysis an Ambion Microbe Express kit (Invitrogen) was used to enrich mRNA. cDNA was synthesized using a SuperScript First Strand Synthesis kit (Invitrogen). qPCR was performed using Luna Universal qPCR Master Mix (New England Biolabs) with 1:5 diluted cDNA in technical duplicate in a StepOne Plus qPCR instrument (Applied Biosystems). A standard curve made from plasmid encoding the gene of interest or a purified PCR product was used to enumerate gene copies in each sample. A no-RT control sample was used to ensure no DNA contamination. Primers were designed using the Integrated DNA Technologies Primer Quest tool.

### Electrophoretic mobility shift assays (EMSAs)

Unlabeled DNA fragments (200-250 bp) encompassing putative promoter regions were amplified from purified *C. difficile* 630Δ*erm* genomic DNA, purified with a GeneJet gel extraction kit (Thermo Fisher), and used as cold competitors or as templates for PCR with IRDye800-conjugated primers (Integrated DNA Technologies). The labeled DNA fragments were purified with a GeneJet PCR purification kit (Thermo Fisher). 20 fmol labeled DNA (or 20 fmol labeled with 1,000 fmol unlabeled for cold competitor control) was mixed with purified BapR and DMSO or 1 μM LCA in binding buffer (20 mM Tris-HCl pH 8, 10 mM KCl, 2 mM MgCl_2_, 0.5 mM EDTA, 1 mM DTT, 0.05% Nonidet-P40, 12% v/v glycerol, 25 μg/mL salmon sperm DNA) for 20 minutes at room temperature and run on a 8% native polyacrylamide gel at 225V at 4°C in the dark. DNA was visualized using an Odyssey CLx imager (LI-COR).

### Data visualization and statistics

All graphs were generated using Prism 9 software (GraphPad). Chemical structures were generated using ChemDraw 20.0 software (Perkin Elmer). Statistical analyses were done using Prism 9 software (GraphPad).

