## Supplementary Figures for "Identification of a bile acid-binding transcription factor in *Clostridioides difficile* using chemical proteomics"

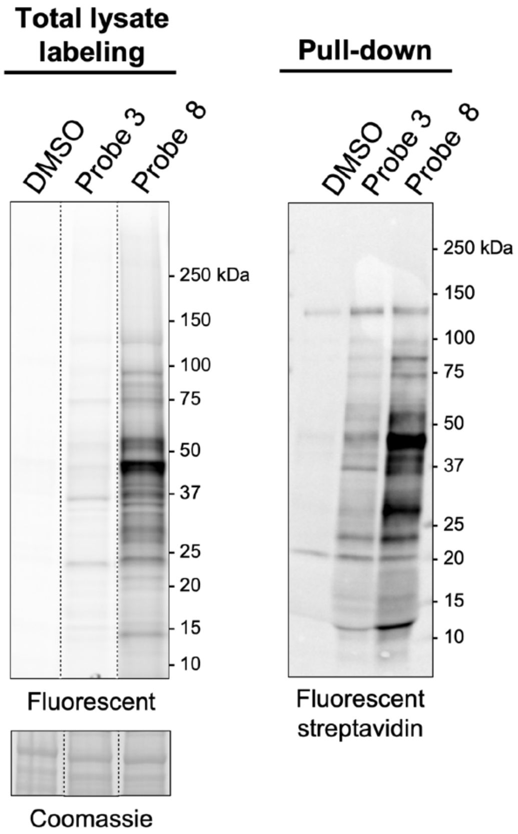


Supplementary Figure 1. Qualitative labeling of *C. difficile* proteins with bile acid probes and isolation of these proteins for proteomics. Fluorescent SDS-PAGE of probe-bound proteins in total lysate as in Figure 1D with Probe 3 included (left) and isolation of proteins with biotin-conjugated bile acid probes using streptavidin beads as the input for the proteomics screen (right); 10 μM probe treatment for 1 hour during log phase, gels are representative of 3 biological replicates.

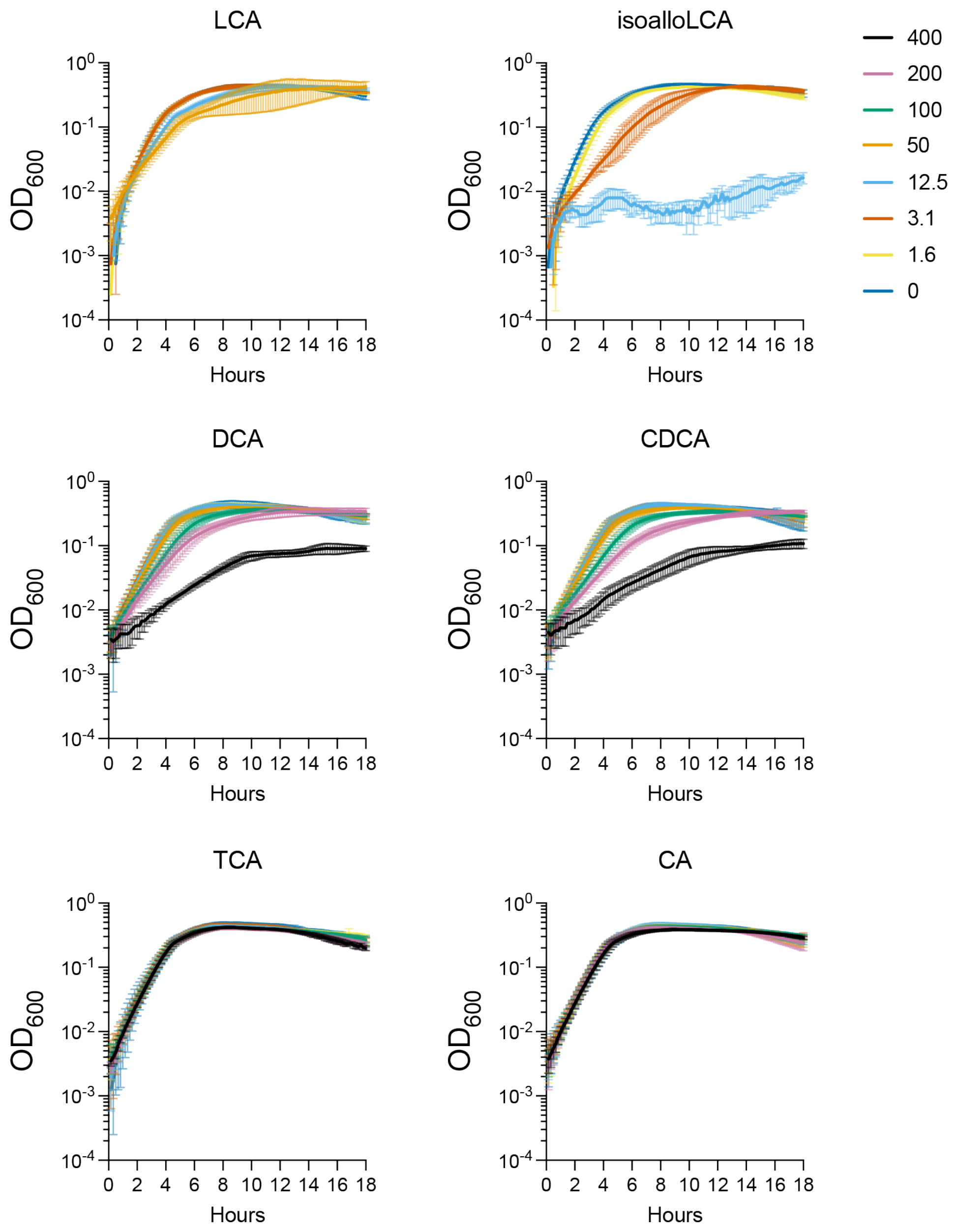


**Supplementary Figure 2. Growth curves of *C. difficile* with bile acids.** Growth measured by $OD_{600}$ across a range of bile acid concentrations; % growth inhibition shown in Figure 2F was calculated from the $OD_{600}$ at 5 hours, n = 4 in two independent experiments.

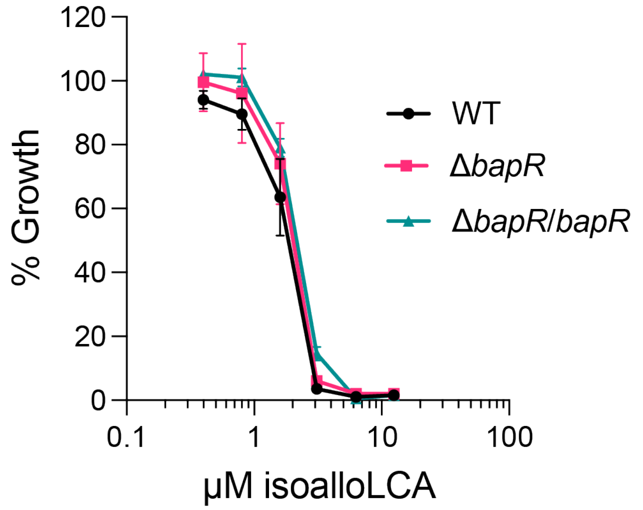


**Supplementary Figure 3. BapR does not affect inhibition of *C. difficile* growth by isoalloLCA.** Growth measured by $OD_{600}$ across a range of isoalloLCA concentrations; % growth inhibition shown in Figure 2F was calculated from the $OD_{600}$ at 5 hours, n = 2 biological replicates in one experiment.

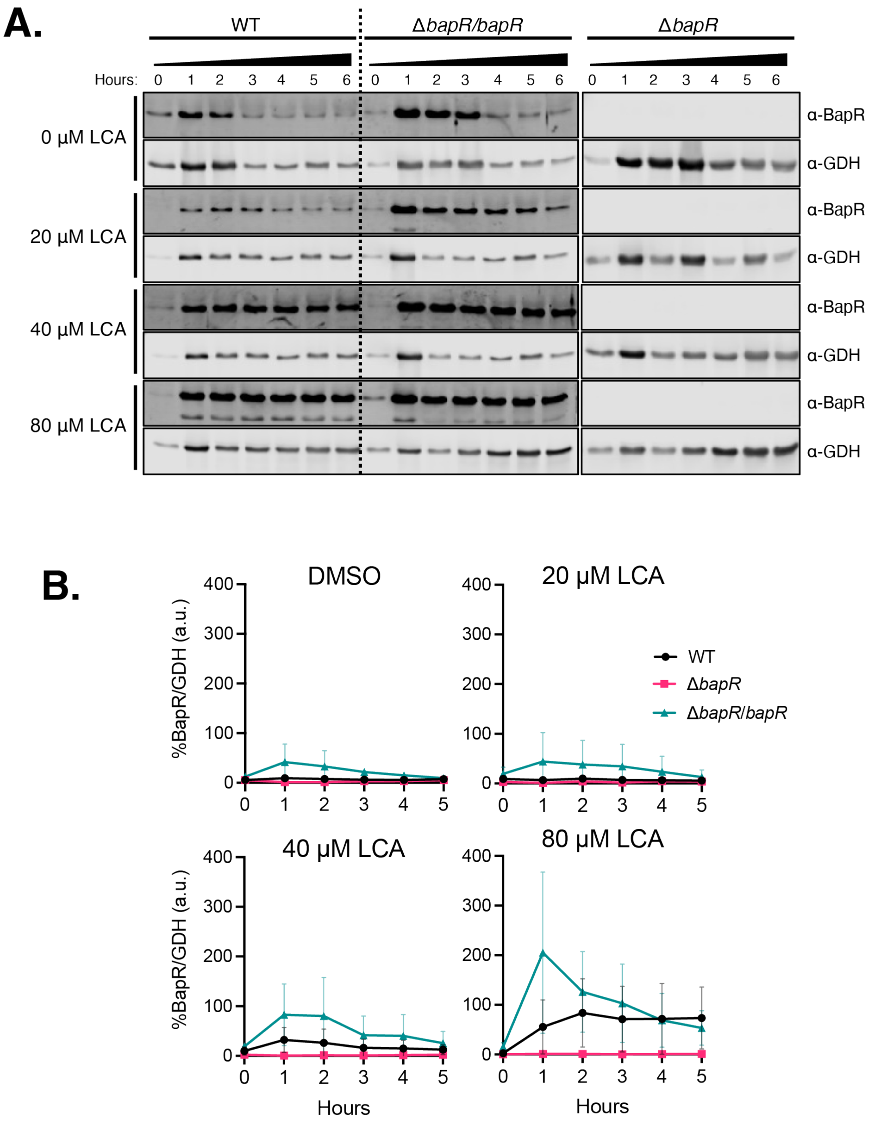


**Supplementary Figure 3. Time course of BapR stabilization by LCA.** (**A**) LCA was added to log-phase *C. difficile* cultures at the indicated concentrations and samples were taken at the indicated timepoints for Western blotting; glutamate dehydrogenase (GDH) serves as a loading control and blots are representative of 3 biological replicates. (**B**) Quantification of the blots in (A); n = 3.

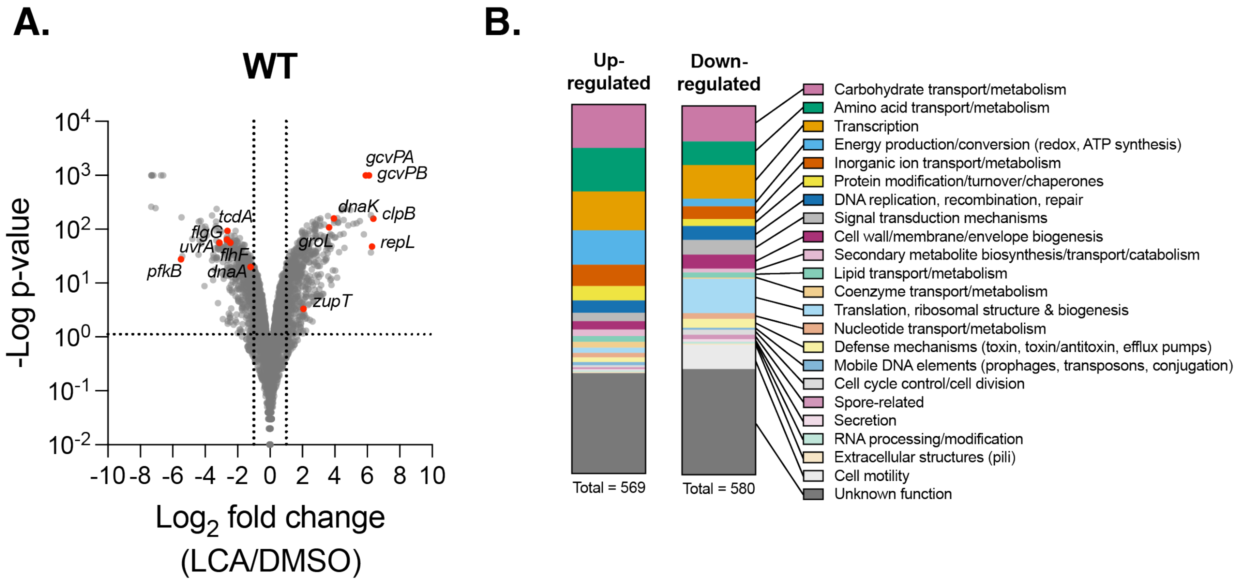


Supplementary Figure 4. Global changes in WT *C. difficile* gene expression in response to LCA. (A) RNA-seq analysis of WT and Δ*bapR C. difficile* after 1-hour treatment with DMSO vehicle or 20 μM LCA during log-phase; dashed lines indicate significance cutoffs at $p < 0.05$ and fold change > 2, n = 3. (B) Classification of differentially expressed genes from (A) by COG category.

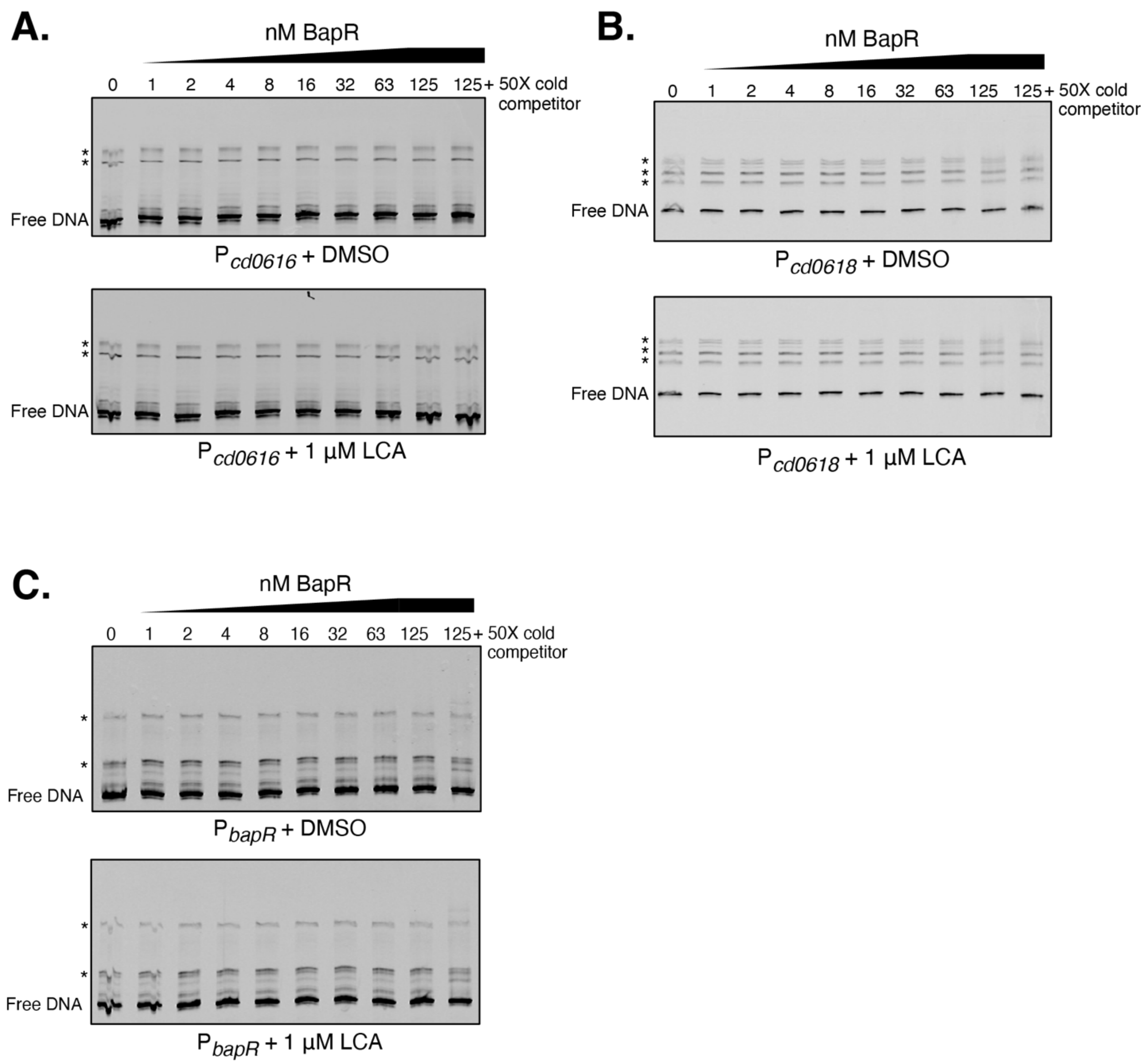


**Supplementary Figure 5. BapR indirectly regulates *cd0618* and *cd0616* and does not bind its own promoter.** (**A**) Electrophoretic mobility shift assay with purified BapR and a 250 bp DNA fragment comprising the region immediately upstream of *cd0616* as a putative promoter; 20 fmol 5' IRDye800-labeled promoter fragment per lane, the last lane contains 20 fmol labeled DNA and 1,000 fmol of the same DNA fragment lacking the fluorescent label as a cold competitor. Gel is representative of 2-3 replicates. (**B**) Assay as in (A) with a 250 bp DNA fragment comprising the region immediately upstream of *cd0618*. (**C**) Assay as in (A) with a 203 bp DNA fragment comprising the region immediately upstream of *bapR*.
